# Extracellular matrix-enriched human induced chondrocyte organoids as living building blocks for scalable cartilage tissue engineering

**DOI:** 10.64898/2026.11.23.768121

**Authors:** Daphne M.A. Menssen, Giorgia Mazzini, Florencia Abinzano, Marina van Doeselaar, Yolande F.M. Ramos, Sebastien J.P. Callens, Ingrid Meulenbelt, Keita Ito

## Abstract

Articular cartilage defects are common and often develop into painful and debilitating osteoarthritis due to the low innate healing capacity of cartilage. However, current regenerative treatments are limited as they depend on large autologous cell numbers and poor scalability. Here, a novel approach for rapid and plentiful cartilage organoid production is introduced for modular and scalable cartilage tissue engineering, by combining an off-the-shelf cell source with a natural matrix supplement in spinner flask cultures. First, human induced chondroprogenitor cell (hiCPC) colonies are matured towards human induced chondrocytes (hiCHO) aggregates in spinner flasks. Although maturation is achieved, resulting hiCHO aggregates show some phenotypic heterogeneity between clones. Nevertheless, subsequent dissociation and culture with a tissue-derived matrix supplement results in rapid and homogeneous organoid self-assembly, generating large numbers of hiCHO-derived cartilage organoids. This matrix-supplemented culture generates consistent pericellular and extracellular matrix-rich organoids without detected off-target cells, limiting the need of labor-intensive selections steps and enhancing scalability. Impressively, 1 million hiCHOs produce over 27,000 organoids within two weeks, with an approximately 30-fold increase in cell number. Finally, hiCHO-derived organoids fuse together to assemble large cartilage constructs, demonstrating their potential as scalable, off-the-shelf building blocks and addressing an unmet need in regenerative cartilage tissue engineering.

## Introduction

Articular cartilage defects in the knee are commonly seen in the clinic, where approximately 60% of all patients under-going arthroscopy, show some degree of cartilage damage.^[1]^ Cartilage defects also occur in young and active patients, often as a consequence of traumatic injury.^[2]^ Articular cartilage has a limited regenerative capacity, due to the absence of blood vessels and lymphatics, as well as its low cellularity.^[3,4]^ Consequently, untreated cartilage defects may predispose the joint into a progressive and negative cascade towards osteoarthritis (OA), a degenerative joint disease that affects hundreds of millions worldwide and causes chronic pain and disability. Given this high burden that OA imposes on society, early interventions are needed to prevent or delay the progression of articular cartilage defects into OA.^[5]^

Tissue engineered cartilage offers a promising solution for the regenerative treatment of cartilage defects. Today, the clinical gold standard for the cell-based treatment of cartilage defects is autologous chondrocyte implantation (ACI).^[6]^ ACI involves harvesting a biopsy from a non-load bearing region of the knee, from which human primary articular chondrocytes (hPACs) are isolated.^[7,8]^ In earlier generations of ACI, expanded hPACs were directly implanted into the defect and covered with a periosteal flap or collagen membrane, while later generations introduced cell-seeded 3D scaffolds such as hydrogels or sponges.^[9]^ More recently, expanded hPACs have been cultured into spheroids, embedded in their own produced but limited extracellular matrix (ECM), and are sparsely seeded on the defect bottom from which they slowly grow *in situ*.^[10,11]^ Although the results of this treatment are encouraging,^[11,12]^ there are important scaling limitations. Taking biopsies of healthy cartilage to obtain autologous hPACs can cause donor site morbidity.^[12]^ In addition, these two-step procedures are six to eight weeks apart to allow for cell expansion and some ECM formation,^[13]^ resulting in a large patient burden with high associated costs. Furthermore, expanding hPACs in order to get a clinically relevant number of autologous cells is limited as passaging induces dedifferentiation towards a fibroblastic phenotype.^[14]^ As a result, passaging must be balanced against cell yield to limit the possible formation of type I collagen-rich fibrocartilage, which is mechanically inferior to native hyaline cartilage and can compromise tissue quality over time.^[15,16]^ As such, there is a clear unmet need for more cost-effective regenerative therapies that reduce the dependency on autologous cells and offer scalable and reliable cartilage production.^[6]^ A promising strategy is to combine human induced pluripotent stem cell (hiPSC)-derived cells with a natural tissue-derived matrix additive to enable the rapid production of large volumes of cartilage tissue.

Recently, hiPSCs have gained significant attention in cartilage tissue engineering, showing promising results both *in vitro* and *in vivo*.^[1–5]^ Human induced chondrocytes (hiCHOs) can be derived from hiPSCs via different intermediate cell states, such as human induced chondroprogenitor cells (hiCPCs) or human induced mesenchymal stem cells (hiMSCs). Of these, cartilage tissue made with hiCPCs has been shown to resemble cartilage made with hPACs more closely.^[3,5]^ This makes hiCPCs an attractive expandable progenitor population with the potential to serve as an off-the-shelf cell source for future cartilage tissue engineering.

Nevertheless, culture times necessary for these cells to produce a sufficient amount of ECM remain considerable, and shortening this phase would further improve scalability. We have previously shown that hPACs cultured in agitated suspension cultures, such as spinner flasks, can self-assemble into cartilage organoids when supplemented with a natural cartilage-like matrix.^[6,7]^ These organoids exhibited cartilage specific properties, including distinguished pericellular matrix and interterritorial matrix with sparsely populated chondrocytes. The matrix supplement, notochordal cell-derived matrix (NCM), is rich in type II collagen and glycosaminoglycans (GAGs), and it has been shown to stimulate chondrocyte proliferation without dedifferentiation, providing the chondrocytes with plentiful cartilage-like ECM directly at the start of the culture.^[6–8]^ Together, these effects enable the quick production of a large volume of cartilage tissue. However, the ability to combine this matrix supplement with hiCPCs for off-the-shelf organoid production has not yet been investigated.

Therefore, the aim of this paper was to combine these two concepts of using an abundant, off-the-shelf cell source and the natural matrix supplement to rapidly create many cartilaginous building blocks for scalable cartilage tissue engineering. The hiCPC colonies were matured into hiCHO aggregates using scalable spinner flasks, as dynamic maturation is less labor intensive than the conventional static maturation and improves matrix deposition.^[1]^ Resulting aggregates were dissociated, and single hiCHOs were cultured in spinner flasks and supplemented with the cartilaginous matrix additive to generate a large volume of cartilage organoids within two weeks. Subsequently, the organoids were characterized and compared with extended matured hiCHO aggregates, a common type of cartilage microtissue and building block. Finally, as a proof-of-concept, the potential of these hiCHO-derived organoids to serve as building blocks for fusing into larger cartilage tissues was evaluated (Figure 1).

**Figure 1.**
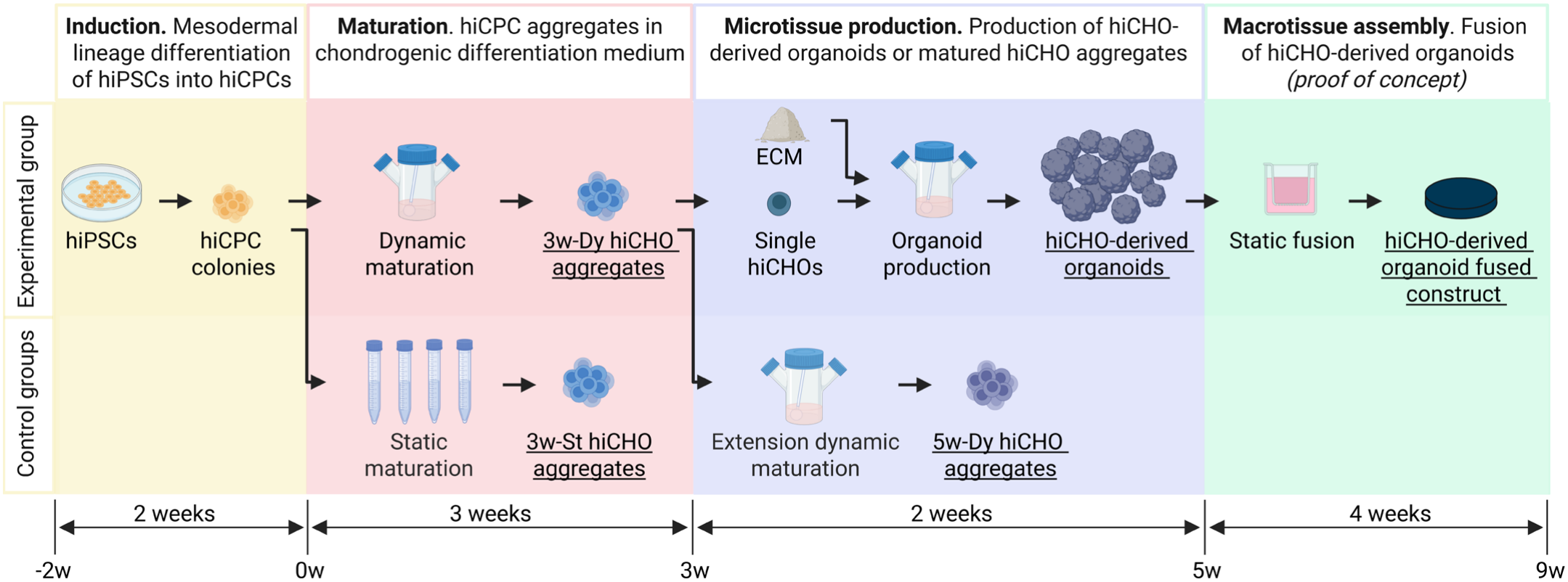
Schematic overview of the experimental setup. hiPSCs (human induced pluripotent stem cells) were induced towards hiCPCs (human induced chondroprogenitor cells). Picked hiCPC colonies were dynamically matured into human induced chondrocyte aggregates for three weeks (3w-Dy hiCHO aggregates) and compared to conventional statically matured hiCHO aggregates (3w-St hiCHO aggregates). Resulting single hiCHOs were supplemented with a cartilaginous ECM (extracellular matrix) in spinner flasks to obtain hiCHO-derived organoids. They were compared to extended dynamically matured hiCHO aggregates (5w-Dy hiCHO aggregates). Finally, hiCHO-derived organoids were used as building blocks to fuse into larger cartilage constructs.

## Results

### Successful dynamic maturation of hiCHO aggregates

We first set out to establish a short and highly scalable spinner flask-based protocol for hiCPC maturation. Previous studies demonstrated that hiCPC colonies maturation towards hiCHO aggregates is possible in both static and dynamic suspension cultures.^[1,2]^ Therefore, scalable spinner flasks were used to dynamically mature hiCPC colonies into hiCHO aggregates for three weeks (3w-Dy) and they were compared to statically matured hiCHO aggregates (3w-St). The three-week maturation period of hiCPC colonies into hiCHO aggregates in spinner flasks was successful, as indicated by the presence of GAGs in most of the 3w-Dy hiCHO aggregates (Figure 2A). Maturation in spinner flasks increased the amount of sulfated GAGs over DNA significantly in the 3w-Dy hiCHO aggregates compared to the 3w-St hiCHO aggregates (Figure 2B). This was accompanied by a lower cell number in the 3w-Dy hiCHO aggregates after maturation, likely due to cells shedding from the periphery of the aggregates in the dynamic culture (Figure S1A and S1B). As indicated by the lack of peripheral GAG production in both maturation conditions (Figure 2A), these peripheral cells are presumably non-chondrogenic.

**Figure 2.**
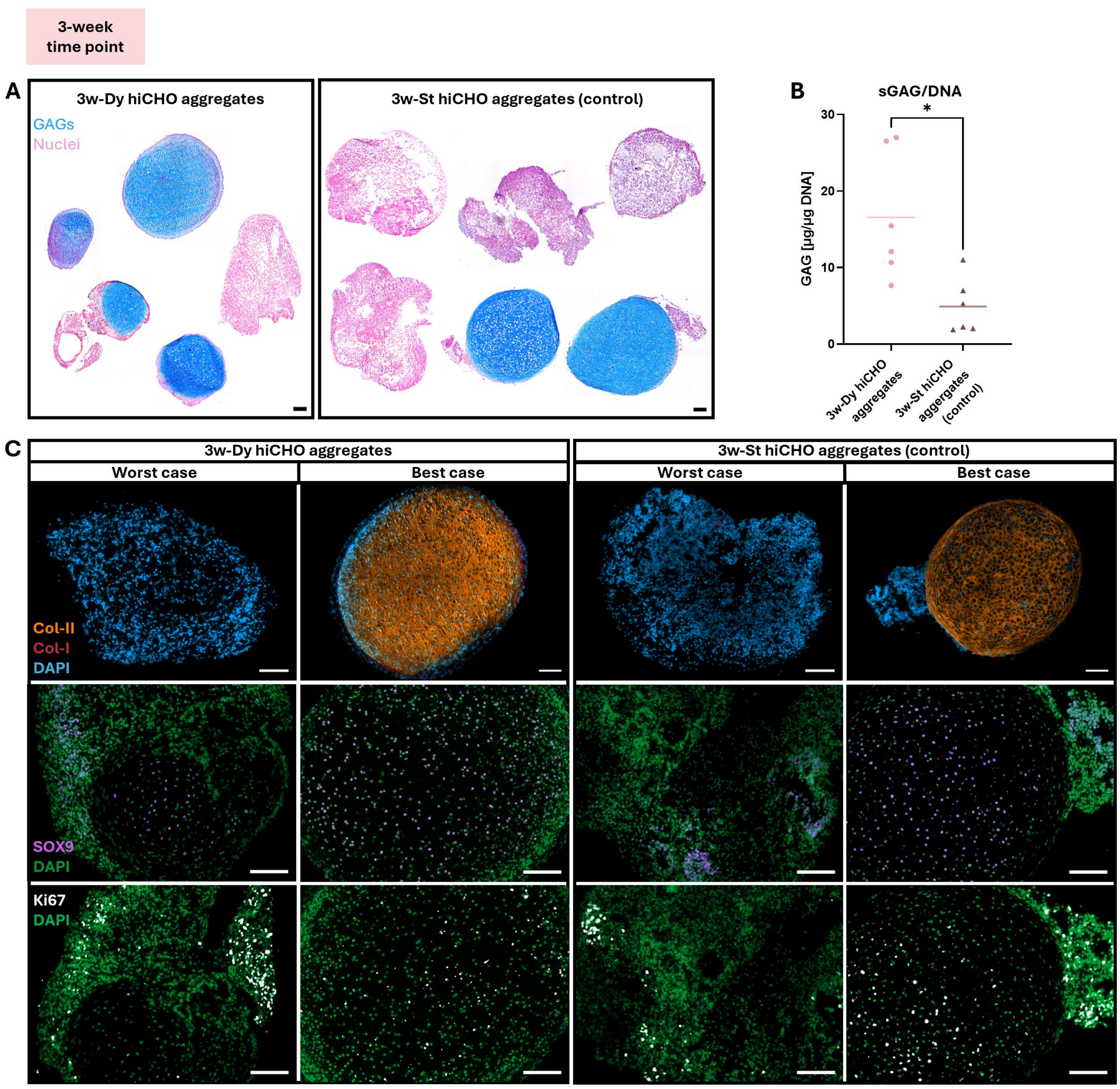
The characterization of hiCHO aggregates matured in either a dynamic (3w-Dy hiCHO aggregates) or static environment (3w-St hiCHO aggregates). Alcian blue staining showing the GAG and cellular distribution in multiple replicates of both groups (A). Quantification of sulfated GAGs normalized against DNA (B). Immunohistochemical analysis for type II collagen, type I collagen, SOX9, and Ki67 of the best and worst 3w-Dy hiCHO aggregates and 3w-St hiCHO aggregates (C). The scale bars are 100 μm. The result of the Welch’s t-test is depicted in the graph, where * indicates P<0.05.

Despite this successful maturation, a large variation between samples was observed for both methods, whereby the GAG content and the type II collagen presence varied substantially between aggregates (Figure 2A and 2C). Prior to spinner flask maturation, hiCPCs naturally formed colonies on 2D culture plastic and therefore contained varying cell numbers. The hiCPC colonies were picked based on their morphology and structural integrity. These observed variation after maturation might be reflected by the differences in the selected hiCPC colonies. The hiCPC colonies were not dissociated and pelleted at a known cell number, as intact hiCPC colonies support a more stable chondrogenic maturation.^[1]^

Aggregates containing type II collagen-rich cores also contained many cells expressing SOX9 (SRY-box transcription factor 9), supporting the chondrogenic phenotype. Interestingly, hiCHO aggregates without any GAGs or type II collagen also contained some SOX9-positive cells (Figure 2C), suggesting delayed maturation in these aggregates. The proliferative capacity, assessed by Ki67 (Kiel 67) staining, appeared comparable between aggregates, suggesting that cell proliferation does not seem to be influenced by the level of chondrogenicity or the maturation method (Figure 2C).

In terms of gene expression, both groups appeared similar after a three-week maturation period (Figure 3). Interestingly, some hiCHO aggregates consistently showed a more chondrogenic phenotype than others, as seen with the high expression of *SOX9, ACAN* (aggrecan), and *COL2A1* (type II collagen) in sample 4 of the 3w-St hiCHO aggregates (Figure 3A, 3B, and 3C). As the trajectory of hiCHO maturation proceeds through a neuroepithelial cell stage, markers associated with alternative cell states were also assessed.^[1–3]^ Specifically, the expression of neurogenic markers, such as *KIF1A* (kinesin family member 1A), has been associated with delayed differentiation upon hiCHO maturation.^[3]^ Indeed, both the 3w-Dy and 3w-St hiCHO aggregates expressed this neuron marker after three weeks in culture at comparable levels to human SH-SY5Y neural cells (Figure 3D). The expression of *COL10A1* (type X collagen) and *COL1A1* (type I collagen), indicating hypertrophic and fibroblastic differentiation respectively, were similar between the two groups, where *COL10A1* expression remained low in the 3w-Dy hiCHO aggregates compared to human chondrocytes derived from knee arthroplasty (Figure 3E and 3F).

**Figure 3.**
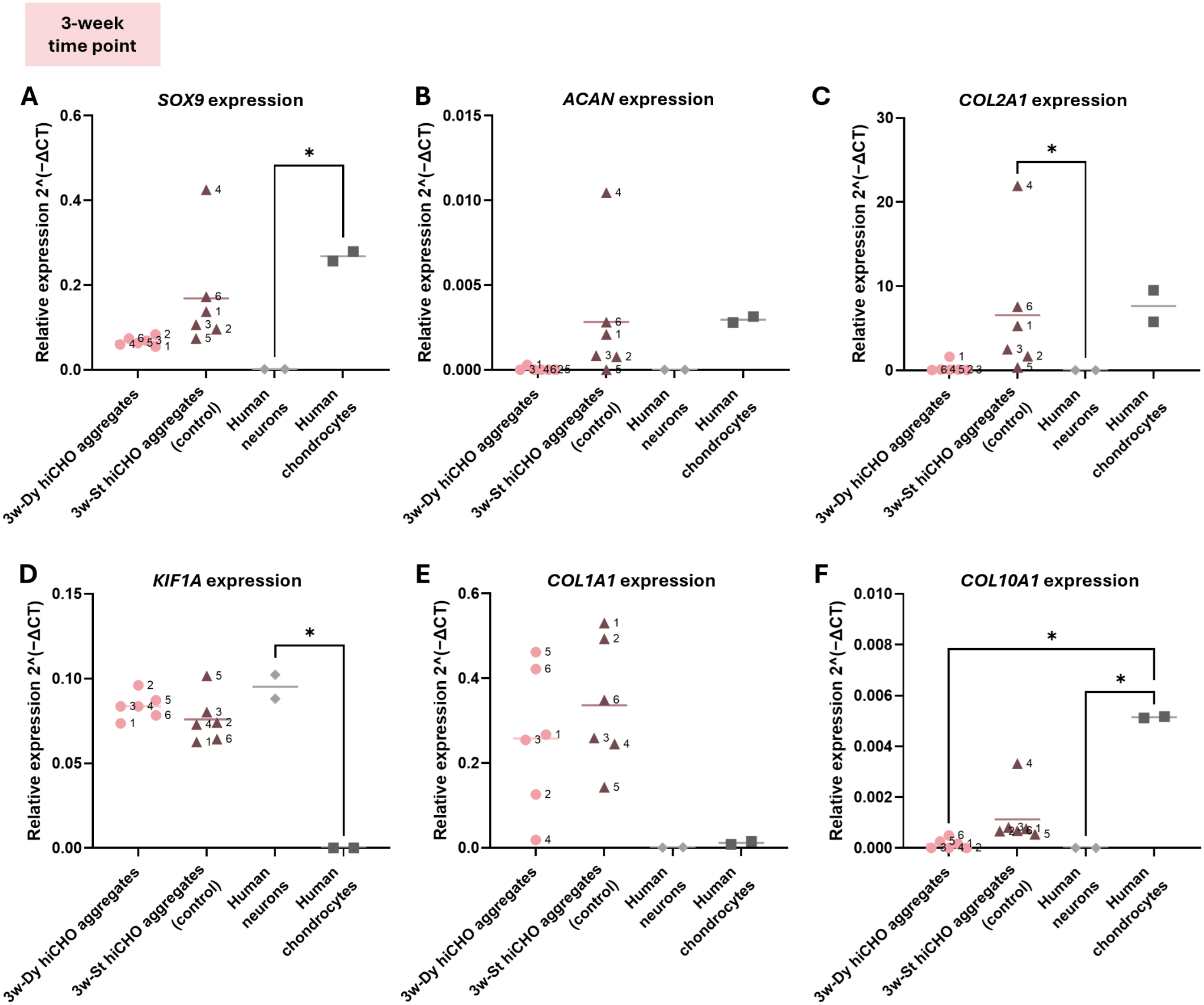
The relative gene expression in 3w-Dy hiCHO aggregates and 3w-St hiCHO aggregates after maturation at the three-week time point. The phenotype of the hiCHOs was compared to human neurons and human chondrocytes. The genes SOX9 (A), ACAN (B), COL2A1 (C), KIF1A (D), COL1A1 (E), and COL10A1 (F) were analyzed. Gene expression was normalized against the house-keeping gene ACTB. The results of Dunn’s multiple comparison tests are depicted in the graphs, where * indicates P<0.05.

### Scalable production of hiCHO-derived cartilage organoids

Following the maturation of the 3w-Dy hiCHO aggregates, the aggregates were dissociated into single cells and cultured in spinner flasks supplemented with the natural matrix supplement, aiming to produce ECM-rich cartilage organoids as modular building blocks for large-scale cartilage tissue engineering. Indeed, our previously established protocols for hPACs could successfully be used to produce an abundant amount of hiCHO-derived cartilage organoids within 14 days, with macroscopic images demonstrating clear consistency across repeated independent cultures (Figure 4A). On average, 1 million hiCHOs resulted in 27,000 organoids with a diameter of 122 ± 74 μm (Figure 4B and 4C). Impressively, this two-week matrix-supplemented culture led to an almost 30-fold increase in cell number (Figure 4D).

**Figure 4.**
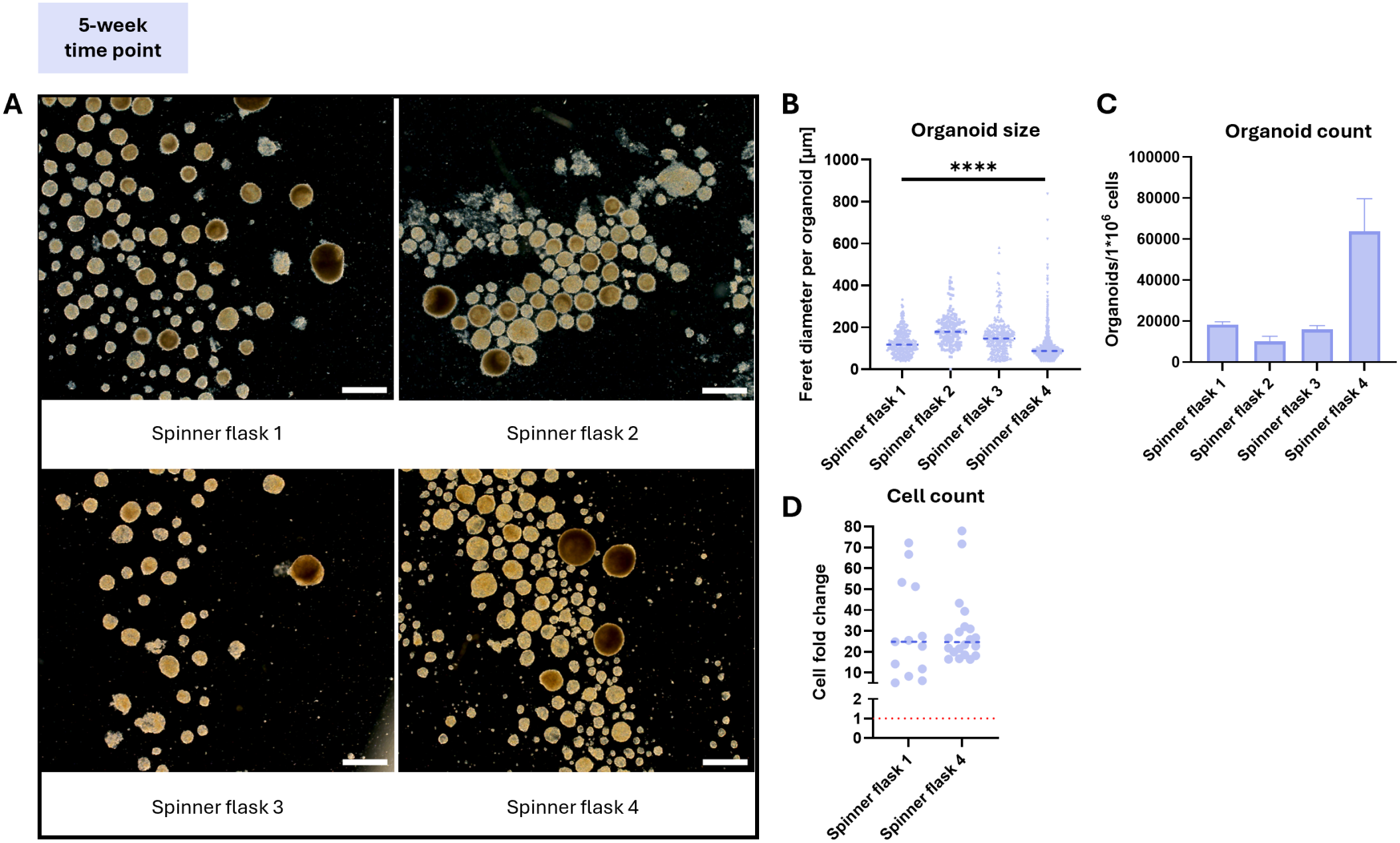
Macroscopic characterization of hiCHO-derived organoids at the five-week time point. Representative microscopy images of the hiCHO-derived organoids formed in independent spinner flasks (A). The Feret diameter distribution after two weeks of matrix-supplemented culture (B). The hiCHO-derived organoid count after two weeks per 1 million hiCHOs seeded on day 0 (N=2) (C). The cell fold change after the hiCHO-derived organoid culture relative to the number of cells initially seeded, which is indicated by the red line (D). The scale bars are 500 μm. The results of the Kruskal-Wallis tests are depicted in the graphs, where **** indicates P<0.0001.

To further assess the effect of matrix supplementation, hiCHO-derived organoids were compared with 5-week dynamically matured (5w-Dy) hiCHO aggregates of the same culture age as the hiCHO-derived organoids, representing a more conventional type of microtissue used for cartilage tissue engineering. The hiCHO-derived organoids showed consistent histological appearance, where the matrix, most likely dominated by the supplemented matrix, filled the center (Figure 5A). The cells mostly occupied the outer periphery, a distribution that likely contributes to future fusion processes.^[4]^ In the 5w-Dy hiCHO aggregates, however, substantial heterogeneity was observed. Nevertheless, all samples showed chondrogenic features as indicated by the cores rich in GAGs, type II collagen, and SOX9-positive cells (Figure 5A and 5D). In contrast, hiCHOs in the organoids did not stain positive for SOX9 (Figure 5D). This was also observed for the proliferation marker Ki67, despite the substantial increase in cell number during the organoid culture (Figure 5D and 4D).

**Figure 5.**
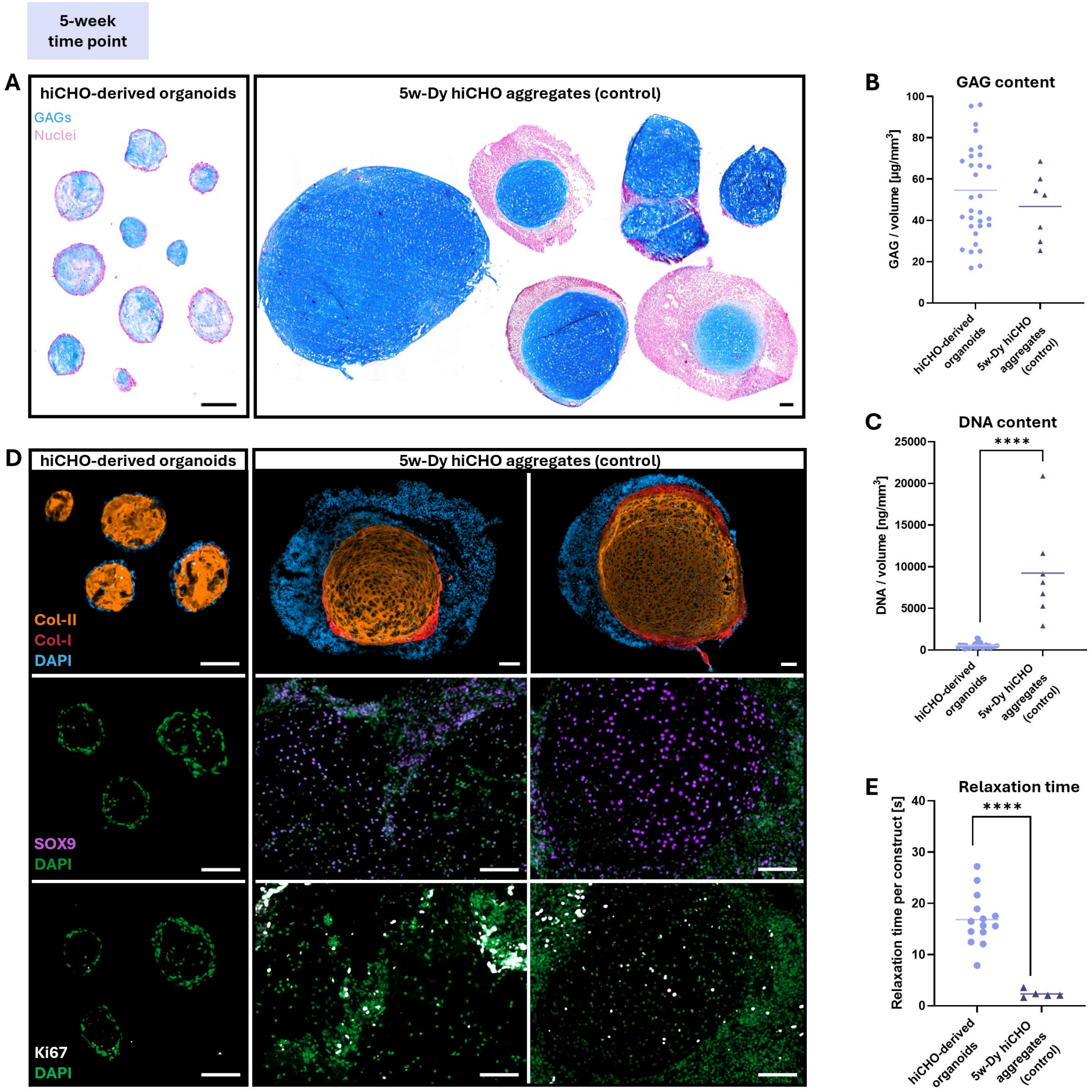
The characterization of hiCHO-derived organoids and 5w-Dy hiCHO aggregates at the five-week time point. Alcian blue staining showing the GAG and cellular distribution in the organoids and aggregates (A). Quantification of sulfated GAGs (B) and DNA normalized against volume of the microtissue (C). Immunohistochemical analysis of type II collagen, type I collagen, SOX9, and Ki67 in hiCHO-derived organoids and 5w-Dy hiCHO aggregates (D). The relaxation time of the microtissues (E). The scale bars are 100 μm. The results of the Welch’s and Mann-Whitney t-tests are depicted in the graphs, where **** indicates P<0.0001. The scale bars are 100 μm.

Interestingly, the two-week-old hiCHO-derived organoids achieved comparable sulfated GAG levels per unit volume compared to the 5w-Dy hiCHO aggregates (Figure 5B), despite containing significantly fewer cells (Figure 5C). This is for a large part attributed to the cell-driven incorporation of the GAG-rich supplemented matrix into the core of the hiCHO-derived organoids. This increased matrix content without a proportional increase of required cells demonstrates a key advantage for large-scale production.

Next, we investigated the mechanical properties of the two types of microtissues, which is relevant in the context of using these as building blocks for cartilage tissue engineering. Single microtissue-level compression testing revealed significantly longer relaxation times in the hiCHO-derived organoids as opposed to the 5w-Dy hiCHO aggregates (Figure 5E). This is potentially due to the matrix supplemented organoids having a denser and more crosslinked ECM network, compared to the matrix produced by the cells.^[5,6]^

The findings from the histological and immuno-histochemical analysis were supported by the gene expression profiles. The relative expression levels for *SOX9* were near-zero in the hiCHO-derived organoids (Figure 6A), which may reflect a diminished stimulus for the hiCHOs to produce cartilaginous ECM and express *SOX9* in the presence of the supplemented cartilaginous matrix.^[7]^ Contrary, the *SOX9* expression in the 5w-Dy hiCHO aggregates was similar to that of human chondrocytes. The expression of the neural marker *KIF1A* increased in the 5w-Dy hiCHO aggregates compared to the three-week time point, whereas its expression was barely detected in the cells from the hiCHO-derived organoids, indicating a reduced presence of incompletely matured or off-target cells there (Figure 6B). The positive type II collagen and GAG staining in the hiCHO-derived organoids most likely reflects the supplemented matrix as *ACAN* and *COL2A1* were barely expressed (Figure 5A, 5D, 6C, and 6D). This low expression could have been caused by the presence of the dense supplemented matrix, similarly to the reduced *SOX9* expression.^[7]^ Notably, the *COL1A1* expression was elevated in the organoids (Figure 6E), although not significant, while the *COL10A1* expression had an increasing trend in the 5w-Dy hiCHO aggregates (Figure 6F).

**Figure 6.**
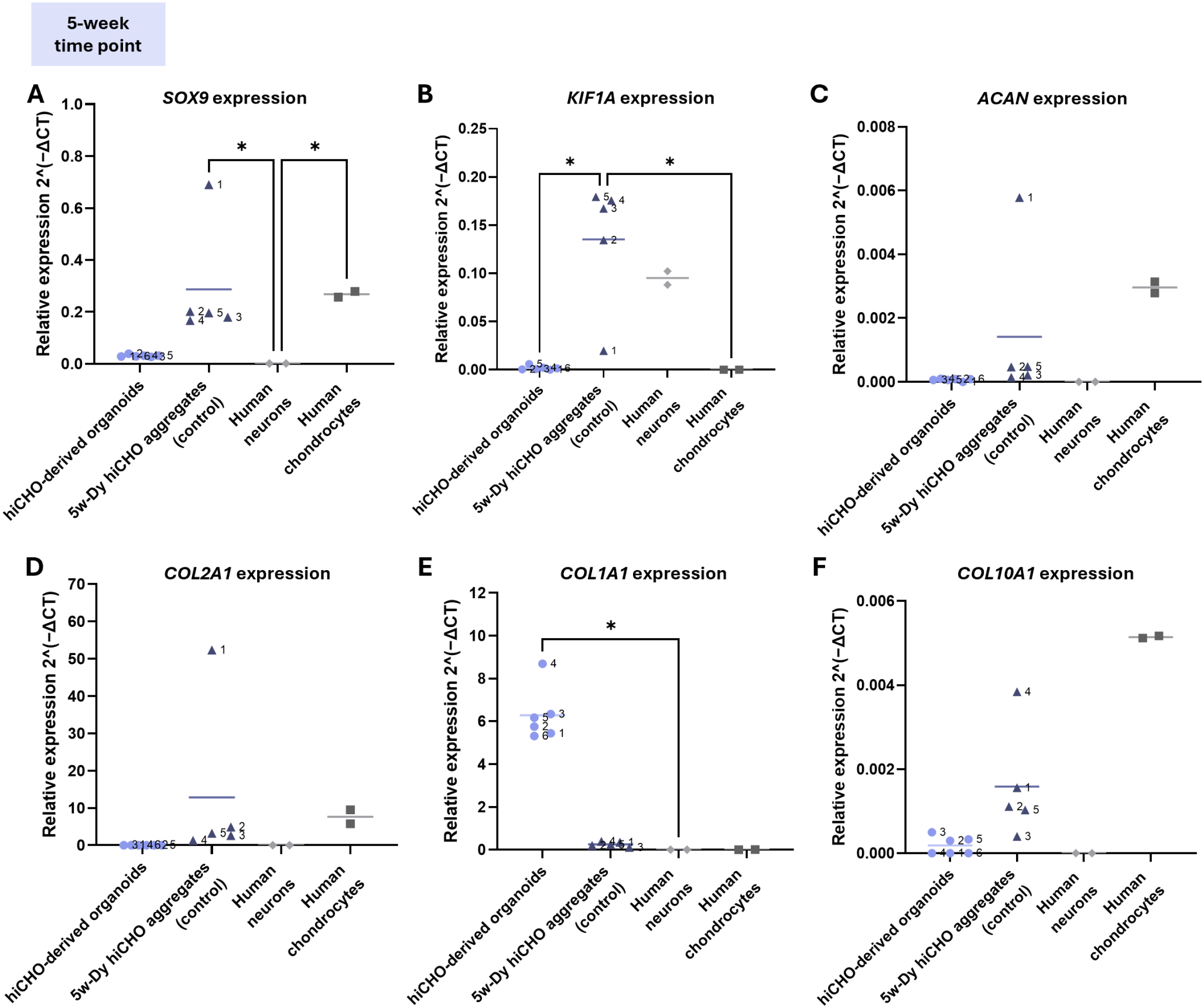
The relative gene expression in hiCHO-derived organoids and 5w-Dy hiCHO aggregates at the five-week time point. The phenotype of the hiCHOs was compared to human neurons and human chondrocytes. The genes SOX9 (A), KIF1A (B), ACAN (C), COL2A1 (D), COL1A1 (E), and COL10A1 (F) were analyzed. Gene expression was normalized against the house-keeping gene ACTB. The results of Dunn’s multiple comparison tests are depicted in the graphs, where * indicates P<0.05.

### The hiCHO-derived cartilage organoids as building blocks for cartilage constructs

As a proof-of-concept exploration for the use of hiCHO-derived organoids as living building blocks for cartilage tissue engineered constructs, the organoids were allowed to fuse in transwells inserts for four weeks (Figure 7A). Organoids successfully fused and grew into a coherent tissue rich in GAGs over time (Figure 7B and 7C). However, we observed tissue compaction at later time-points in all replicates (N=7) (Figure 7D), and the constructs showed a clear expression of type I collagen, especially on the periphery (Figure 7E). The type II collagen observed in the fused constructs most likely originates for a large part from the supplemented matrix in the hiCHO-derived organoids prior to fusion. Interestingly, hiCHOs from the previously quiescent organoids re-expressed SOX9 and showed a slight increase in proliferation, as indicated by Ki67 staining (Figure 7E).

**Figure 7.**
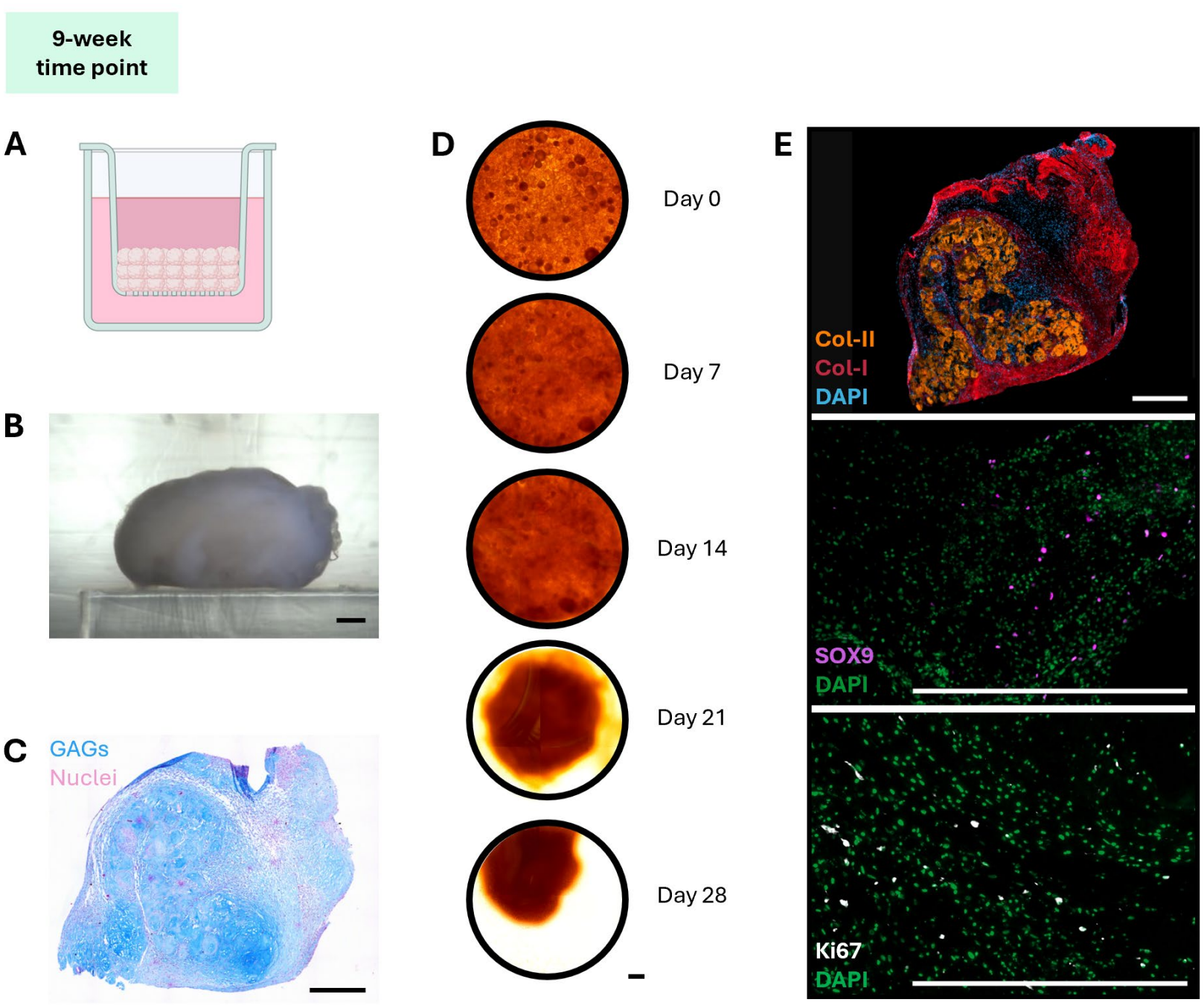
The characterization of hiCHO-derived organoid fused constructs at the nine-week time point. The transwell system used for the fusion of a few layers of hiCHO-derived organoids (A). A macroscopic image of a hiCHO-derived organoid fused construct after a four-week fusion culture (B). Alcian blue staining showed the GAG and cellular distribution in the constructs (C). A timeline showing the top view of the fusion and compaction of the construct over 28 days (D). Immunohistochemical analysis of type II collagen, type I collagen, SOX9, and Ki67 in the hiCHO-derived organoid fused construct (E). The scale bars are 500 μm.

The expression of cartilage-specific markers, including *SOX9, ACAN*, and *COL2A1*, increased during the entire culture, which supports a shift towards a chondrogenic profile over time (Figure 8A, 8B, and 8C). Importantly, the *KIF1A* expression remained low in the fused constructs, which is consistent with the low expression observed after the matrix supplemented organoid culture (Figure 8D). However, *COL1A1* expression levels increased, consistent with immunohistochemical analysis (Figure 8E). Type I collagen production can be associated with fibrocartilage and contractile chondrocytes, and it is a common challenge within cartilage tissue engineering.^[8–10]^ However, it can also indicate the immature state of the engineered tissue, as the early stages of cartilage development is characterized by type I collagen which is slowly remodeled into type II collagen.^[11]^ This immaturity is further supported by the low *COMP* (Cartilage Oligomeric Matrix Protein) and *CHAD* (chondroadherin) expression compared to human chondrocytes, which are associated with ECM organization (Figure 8F and 8G).^[12,13]^ As expected, hypertrophic differentiation was not observed, as the expression of *MGP* (Matrix Gla Protein) and *COL10A1* remained low compared to human chondrocytes (Figure 8H and 8I).^[13–15]^

**Figure 8.**
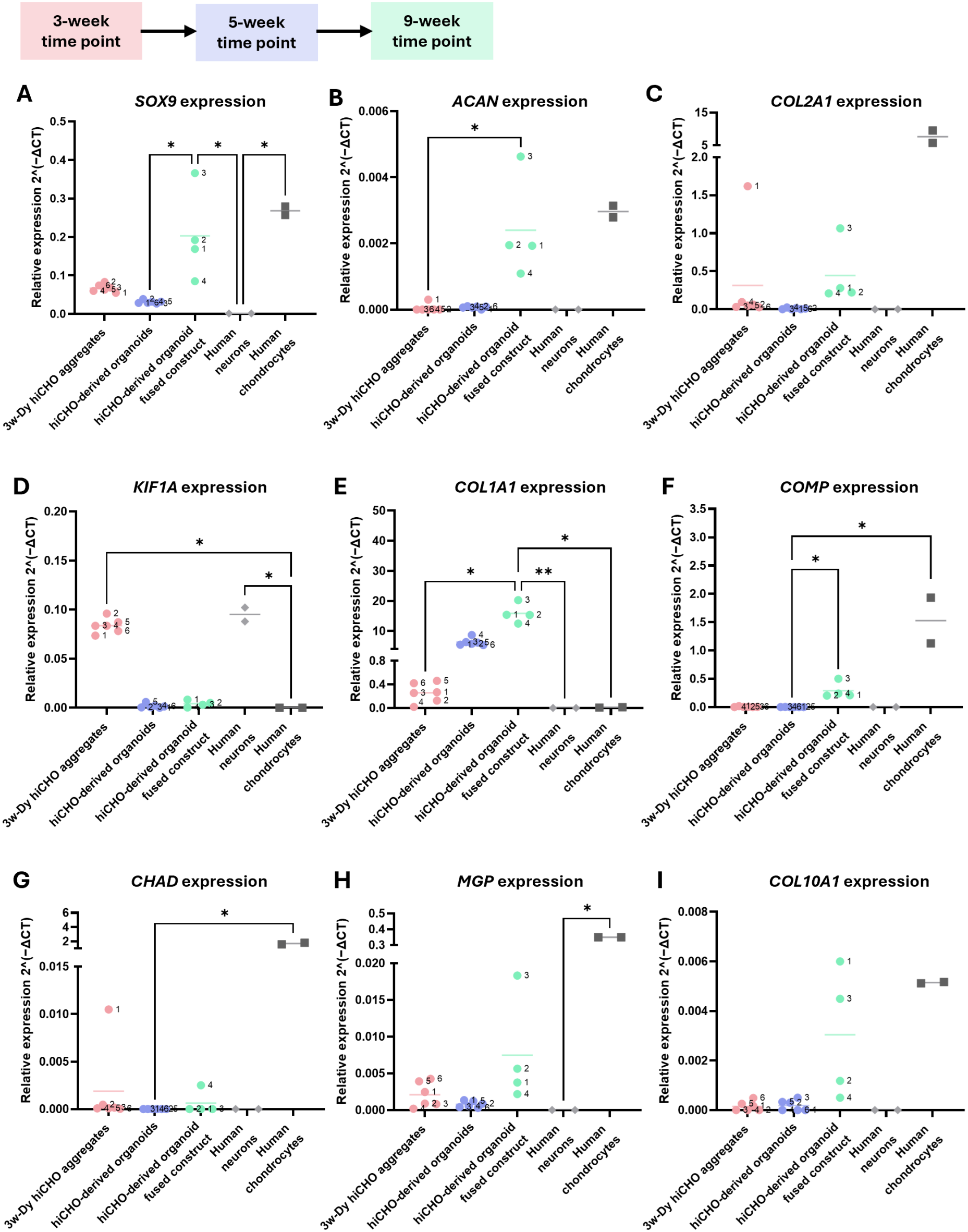
The relative gene expression in the 3w-Dy hiCHO aggregates, the hiCHO-derived organoids, and the hiCHO-derived organoid fused constructs. The phenotype of the hiCHOs was compared to human neurons and human chondrocytes. The genes SOX9 (A), ACAN (B), COL2A1 (C), KIF1A (D), COL1A1 (E), COMP (F), CHAD (G), MGP (H), and COL10A1 (I) were analyzed. Gene expression was normalized against the house-keeping gene ACTB. The results of Dunn’s multiple comparison tests are depicted in the graphs, where * indicates P<0.05.

## Discussion

The aim of this paper was to develop a novel approach for scalable cartilage tissue engineering by producing a large number of ECM-enriched building blocks from relatively low numbers of hiCPCs, a cell source with off-the-shelf potential. To achieve this high degree of scalability, hiCPC colonies were first matured dynamically towards hiCHO aggregates using spinner flasks. After this short maturation stage, the resulting hiCHO aggregates were dissociated into single cells and supplemented with a tissue-derived matrix additive in spinner flasks to promote the spontaneous and large-scale self-assembly of hiCHO-derived organoids. These organoids exhibited key ECM characteristics of native cartilage, thereby indicating strong potential as building blocks for cartilage tissue engineering.

Chondrogenic maturation in spinner flasks generally supported hiCPC to hiCHO differentiation. Despite four-to six-week maturation periods being more common for induced chondrocytes, the three-week maturation resulted in chondrogenic aggregates.^[13,15,16]^ Interestingly, the neuronal associated marker, *KIF1A*, was expressed in the 3w-St and 3w-Dy hiCHO aggregates at similar levels as human neurons. Recent single nucleus multimodal analysis of hiPSCs to hiCHOs reported that chondrogenic cell fate initially follows a neurogenic developmental program before diverging toward a chondrogenic fate.^[3,17–19]^ Therefore, persistent expression of *KIF1A* may reflect an incomplete chondrogenic commitment rather than a confirmed neuronal off-target population of cells.

Following maturation, the 3w-Dy hiCHO aggregates were dissociated into single cells without selecting for the most chondrogenic aggregates. The resulting hiCHO-derived organoids were consistent in terms of matrix content and gene expression profiles, despite starting with a heterogeneous cell population. Most importantly, the *KIF1A* expression was lost in the organoids. These observations may indicate that the matrix supplement either has a selective effect on chondrogenic cells during organoid self-assembly, or that the cartilage-like environment suppresses the *KIF1A* expression. Regardless of the underlying mechanism, the ability to produce homogeneous hiCHO-derived organoids from an inherently heterogenous hiPSC-derived product indicates that selection steps are not required when using this matrix supplement, increasing scalability.^[2]^

During the matrix-supplemented culture, a large volume of tissue was generated rapidly. This was achieved by providing an ECM to the hiCHOs, reducing the dependence on cell-derived ECM production. On average, 1 million hiCHOs yielded 27,000 organoids within two weeks with a mean diameter of 122 ± 74 μm. Compared to the 5w-Dy hiCHO aggregates, the organoid-based approach enabled the production of microtissues with comparable matrix output while requiring substantially fewer cells. This efficiency also becomes evident when comparing our organoid-based protocol to the spheroid-based ACI method used clinically for cartilage repair, where 3,000 to 200,000 hPACs are required to create one aggregate of 100 to 1,000 μm in diameter.^[20,21]^ In contrast, the organoid-based approach in this study can produce on average 80 to 5,400 small microtissues from the same range of cells. These organoids are attractive building blocks for biofabrication strategies, as their smaller sizes may improve fusion capacity.^[22]^

The cell number increased substantially during the matrix-supplemented culture, consistent with the known proliferative effects of the used matrix supplement.^[23,24]^ Furthermore, the matrix supplement, rich in bioactive factors, is known to stimulate ECM production in nucleus pulposus cells and primary chondrocytes, as well as the chondrogenic phenotype characterized by *SOX9* expression.^[23,25–28]^ However, *SOX9* gene and protein expression were absent in the hiCHO-derived organoids. Even though the prior short three-week maturation phase saves time and costs, the SOX9 absence may suggest that the hiCHOs were not yet mature enough to maintain their chondrogenic phenotype with matrix supplementation only. Therefore, sustaining stimulation with transforming growth factor beta 1 (TGF-β1) during the organoid culture or extending the hiCHO maturation period might be required.^[13,16,29]^

As a proof-of-concept test, hiCHO-derived organoids were fused in transwells to assess their potential as building blocks for larger cartilage constructs. The static fusion, without a scaffold or mechanical stimulation, was successful. Interestingly, hiCHOs, which had previously lost the expression of chondrogenic markers, regained SOX9 gene and protein expression, as well as the increased expression of ECM genes such as ACAN, suggesting a partial recovery of the chondrogenic profile. The presence of type I collagen combined with the low expression of COMP and CHAD could possibly indicate immaturity. Therefore, future studies should focus on optimizing the fusion environment and construct maturation, to more closely resemble conditions found in vivo. Providing the right supporting 3D environment, such as a bulk or granular hydrogel, may help in stabilizing the chondrogenic phenotype.^[48,49]^ The observed compaction of the fused constructs could potentially be related to elevated cellular contractility due to the lack of adhesive boundary interfaces and to a possible fibroblastic phenotype.^[35]^ The incorporation of a scaffold, potentially supported by directional growth factor stimulation, could help guiding the collagen architecture, maintaining the shape of the fused construct, and improving the mechanical properties of the construct.^[50–52]^

Spinner flasks provide a practical and efficient culture platform for the dynamic maturation of hiCHO aggregates and subsequent organoid production (Figure 9). The improved nutrient and oxygen delivery during aggregate maturation reduces medium consumption and associated costs, compared to conventional static cultures.^[35]^ In addition, spinner flasks simplify medium changes, enable the maturation of aggregates simultaneously in one flask, minimize incubator space, and are compatible with automated systems.^[36]^ However, a limitation of this approach is the reduced control over the size and number of organoids produced.^[16]^

**Figure 9.**
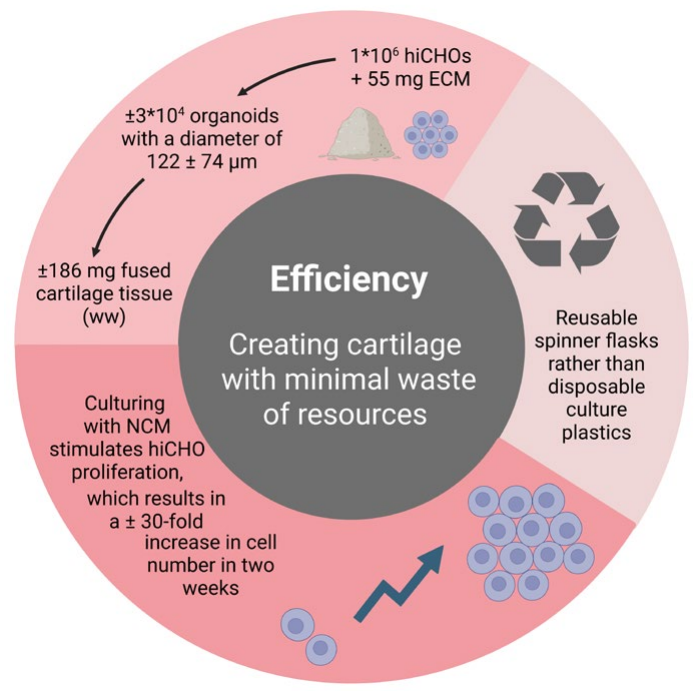
Schematic overview showing the efficiency of making cartilage tissue using the hiCHO-derived organoid-based protocol.

A limitation of this study is that the 5w-Dy hiCHO aggregates were not included in the fusion experiments. While a large volume of hiCHO-derived organoids could be generated, insufficient 5w-Dy hiCHO aggregates could be produced to fill a transwell insert under the current experimental conditions. This comparison would have determined whether fused 5w-Dy hiCHO aggregates showed fewer immature characteristics and less contractile behavior. Furthermore, the porcine-derived matrix supplement used is not fully characterized, and the mechanisms by which self-assembly with chondrocytes is driven, remains unknown. As these limitations are important considerations for clinical translation, we are currently aiming to clarify them to identify a defined natural substitute.

## Conclusion

Current cartilage tissue engineering strategies used in the clinic face scalability challenges. This study introduced an efficient and scalable approach to produce hiCHO-derived organoids. First, hiCPC colonies, with off-the-shelf potential, were matured into hiCHO aggregates and single hiCHOs were subsequently cultured in a matrix-supplemented dynamic suspension culture. The dissociation of hiCHO aggregates followed by self-assembly with a natural matrix supplement resulted in the fast production of a high cartilage organoid volume, with an approximately 30-fold increase in cell number. The combination of an hiPSC-derived cell source and matrix supplementation diminished the need for autologous cells and minimized the time required for cell-driven ECM production, respectively. Importantly, the matrix-supplemented organoid culture yielded a homogeneous product, despite starting from a heterogeneous hiCHO cell pool. Therefore, labor-intensive selections steps are not required, enhancing the efficiency and scalability of the strategy. Finally, the hiCHO-derived organoids fused into larger cartilage constructs, demonstrating their potential as off-the-shelf and scalable building blocks for cartilage tissue engineering.

## Acknowledgements

Microcopy was performed at the LCTE Microscopy Facility, TU/e. The monoclonal antibody for type II collagen (II-II6B3), developed by T. F. Linsenmayer, was obtained from the Developmental Studies Hybridoma Bank, created by the NICHD of the NIH and maintained at The University of Iowa, Department of Biology, Iowa City, IA 52242.

## Author contributions

DMAM: Conceptualization, Investigation, Writing - original draft, Methodology, Visualization, Writing - review & editing, Formal analysis, Project administration, and Data curation. GM: Conceptualization, Investigation, Writing - original draft, Writing - review & editing, Methodology, and Formal analysis. FA: Conceptualization, Writing - review & editing, Supervision, and Methodology. MVD: Investigation, Methodology, Formal analysis, and Writing - review & editing. YFMR: Conceptualization, Writing - review & editing, Supervision, Methodology, and Formal analysis. SJPC: Conceptualization, Project administration, Supervision, Writing - review & editing, Methodology, Visualization, and Formal analysis. IM: Conceptualization, Methodology, Writing - review & editing, Supervision, Funding acquisition, and Formal analysis. KI: Conceptualization, Methodology, Visualization, Writing - review & editing, Project administration, Supervision, Funding acquisition, and Formal analysis.

## Data availability statement

Data can be made available upon request.

## Materials and Methods

### Culture

#### Sample description and ethics approval

In this study, the hiPSC line LUMC0099iCTRL04 was used. Cells were generated after consensus from skin fibroblasts of a female donor by the LUMC iPSC Core facility and registered at the Human Pluripotent Stem Cell Registry under the number P13.080. Cells were characterized according to the pluripotent potential and spontaneous differentiation capacity by the iPSC Core Facility.^[55]^

#### hiPSCs culture and maintenance

The hiPSCs were maintained in mTesR Plus medium (STEMCELL Technologies) on VitronectinXF-coated plates (STEMCELL Technologies) at 37°C in 5% CO_2_. Medium was refreshed every two days and the cells were passaged using Gentle Cell Dissociation Reagent (STEMCELL Technologies) at approximately 80% confluency. Mycoplasma tests were performed on a regular basis.

#### Stepwise hiPSC induction towards hiCPCs

Generation of hiCPCs was based on previously described protocols.^[26,41]^ When hiPSCs reached 60% confluency, mTesR Plus medium was switched to mesodermal differentiation (MD) medium composed of IMDM Gluta-MAX (IMDM; Thermo Fisher Scientific) and Ham’s F12 Nutrient Mix (F12; Sigma-Aldrich) supplemented with chemically defined lipid concentrate (1%, Gibco), insulin/human transferrin/selenous-plus premix (ITS+ premix; 1%, Corning), penicillin-streptomycin (P/S; 0.5%, Gibco), and 1-thioglycerol (450 μM, Sigma-Aldrich). On day 0, hiPSCs were washed with wash medium (IMDM/F12 and 0.5% P/S) and cultured in MD medium supplemented with activin A (30 ng mL^-1^, Miltenyi Biotec), CHIR99021 (CHIR; 4 μM, Tocris), and human fibroblast growth factor-2 (FGF-2; 20 ng mL^-1^, R&D Systems) for 24 hours. On day 1, cells were washed with wash medium and cultured in MD medium supplemented with SB-505124 (2 μM, Tocris), CHIR (3 μM), FGF-2 (20 ng mL^-1^), and dorsomorphin (4 μM, Tocris) for 24 hours. On day 2, cells were washed again and cultured in MD medium supplemented with SB-505124 (2 μM), dorsomorphin (4 μM), C59 (1 μM, Tocris), and PD173074 (500 nM, Tocris) for 24 hours. From days 3 to 5, cells were washed daily with wash medium and cultured in MD medium supplemented with purmorphamine (2 μM, Tocris) and C59 (1 μM). From days 6 to 14, cells were washed briefly with wash medium daily and cultured in MD medium supplemented with human bone morphogenetic protein 4 (BMP-4; 20 ng mL^-1^, Miltenyi Biotec).

#### hiCPC maturation towards hiCHOs

Following the two-week induction of hiPSCs to hiCPCs, the resulting hiCPC colonies were picked from the culture plastic and collected in custom made spinner flasks in chondrogenic differentiation (CD) medium.^[22,26,40]^ CD medium consisted of DMEMF-12 (10565018; Gibco), ITS+ premix (1%), MEM non-essential ammino acids (NEAA; 1%, Gibco), dexamethasone (0.1 μM, Sigma-Aldrich), P/S (0.2%), (+)-Sodium L-ascorbate (50 μg mL^-1^, Sigma-Aldrich), 2-Mercaptoethanol (55 μM, Gibco), L-Proline (40 ng mL^-1^, Sigma-Aldrich), and TGF-β1 (10 ng mL^-1^, Preprotech). The hiCPC colonies were allowed to mature for three weeks in the spinner flasks on a magnetic stirring plate (Variomag Biosytem 4; Thermo Fisher Scientific) at 120 rpm controlled by the Biomodul 40B (Thermo Fischer Scientific) in hypoxia (37°C, 5% CO_2_, and 2.5% O_2_). The medium was refreshed twice a week. The resulting hiCHO aggregates were referred to as 3w-Dy hiCHO aggregates. As a control, several picked hiCPC colonies were matured towards hiCHO aggregates statically under the same conditions. Here, medium changes were performed three times a week. The resulting hiCHO aggregates were referred to as 3w-St hiCHO aggregates.

#### hiCHO Organoid Culture

After maturation, the 3w-Dy hiCHO aggregates were dissociated into single hiCHOs, by incubation in pronase (2 mg mL^-1^, Roche), followed by incubation in type I collagenase (1.52 mg mL^-1^, Worthington), both in CD medium for 1 hour at 37°C.^[56]^ The resulting single hiCHOs were used to create cartilage organoids according to a previously published protocol.^[23–25]^ Briefly, the custom made spinner flasks were filled with 50,000 hiCHOs and 0.25 mg NCM per mL organoid medium (high glucose DMEM (419660; Gibco), fetal bovine serum (FBS; 5%, Greiner Bio-One), ITS+ premix (1%), P/S (1%), 4-(2-hydroxyethyl)-1-piperazineethanesulfonic acid (HEPES; 10 mM, Gibco), L-ascorbic acid 2-phosphate (0.2 mM, Sigma-Aldrich), and NEAA (1%)). The porcine NCM was produced as previously described.^[57]^ In short, the nucleus pulposus was harvested under sterile conditions from the intervertebral discs of 12-week-old piglets (obtained as slaughter waste from an abattoir), and lyophilized and pulverized to obtain a fine powder. To improve the survival of the single hiCHOs, CloneR2 (100 μL mL^-1^, STEMCELL Technologies) was added to the spinner flask on the day of seeding. The hiCHO organoids were allowed to grow for two weeks in the spinner flask on a magnetic stirring plate at 120 rpm in hypoxia (37°C, 5% CO_2_, and 2.5% O_2_). The medium was refreshed twice a week and during these medium changes, the NCM concentration increased from 0.25 mg mL^-1^ to 0.5 mg mL^-1^, finally reaching 1 mg mL^-1^.

To evaluate the effect of the matrix supplement on the hiCHOs, several intact hiCHO aggregates were allowed to mature for two more weeks in the spinner flasks. This group was referred to as 5w-Dy hiCHO aggregates.

#### Fusion

To evaluate the potential of the hiCHO-derived organoids as building blocks in cartilage tissue engineering, the organoids were in fused transwells inserts (cellQART 24-Well Cell Culture insert PET; Sterlitech Corporation) coated with rat tail type I collagen (0.125 mg mL^-1^, Corning).^[52]^ After seeding, the organoids in the transwells were centrifuged and allowed to adhere at 37°C for 10 minutes before adding medium. The constructs were submerged in organoid medium supplemented with TGF-β1 (10 ng mL^-1^) and dexamethasone (0.1 μM). They were cultured for 28 days in hypoxia (37°C, 5% CO_2_, and 2.5% O_2_) with medium changes three times a week.

### Analysis

#### Organoid number and size

The size and number of hiCHO-derived organoids was determined at the end of the organoid culture. Two samples (200 μL) were taken from each spinner flask and evaluated using the EVOS XL Core Imaging System (Invitrogen). The number of organoids per sample was counted manually, while the Feret diameter was determined with ImageJ software.^[23]^

#### Biochemical analysis

After culture, the hiCHO aggregates and hiCHO-derived organoids were frozen, lyophilized, and digested for 16 hours at 60°C with a papain digestion buffer (phosphate buffer (100 mM, Sigma-Aldrich), L-cysteine (5 mM, Sigma-Aldrich), ethylenediamine tetraacetic acid (EDTA; 5 mM, Sigma-Aldrich), and papain (130 μg mL^-1^, Sigma-Aldrich)). The sulfated GAG and DNA contents were evaluated using biochemical a 1-9-dimethyl methylene blue (DMMB, pH = 3.0; Sigma-Aldrich) assay^[58]^ and Qubit dsDNA high-sensitivity kit (Invitrogen), respectively. The DNA content was used to determine cell proliferation during the hiCHO-derived organoid culture.

#### Histology

The hiCHO aggregates, hiCHO-derived organoids, and hiCHO-derived organoid fused constructs were fixed overnight at 4°C in formalin (3.7%, Sigma-Aldrich) with eosin (0.1%, Sigma-Aldrich). To simplify the handling of the small tissues, the samples were embedded in sodium alginate (3 w/v%, Sigma-Aldrich) before dehydration in a tissue processor (HistoCore PEARL, Leica Biosystems). Thereafter, the samples were embedded in par-affin (Sigma-Aldrich) and 5 μm sections were made. To visualize the GAGs in the samples, the sections were deparaffinized, rehydrated, and incubated with citrate buffer (10x, DAKO Agilent) to remove the alginate. Subsequently, the sections were stained with alcian blue (1 w/v%, pH=2.5, Sigma-Aldrich) and Nuclear Fast Red (Millipore). Before mounting with Entellan (Sigma-Aldrich), the sections were dehydrated again. Finally, they were imaged with the Leica DMi8 microscope (Leica Microsystems).

#### Immunohistochemistry

To evaluate the matrix components of the tissues and the chondrogenicity of the cells, an immunohistochemistry staining for type II collagen, type I collagen, SOX9, and Ki67 was performed. The sections were again deparaffinized and rehydrated. The antigen retrieval for type I collagen and type II collagen consisted of incubation with pepsin (0.1%, Sigma-Aldrich) in HCl (10 mM, Sigma-Aldrich) at 37°C. Before primary antibody incubation (Table 1), the sections were blocked with normal goat serum (NGS; 5%, Gibco) in phosphate-buffered saline (PBS; Sigma-Aldrich). The antigen retrieval for SOX9 and Ki67 was performed with citrate buffer (1x) at 96°C, and these sections were blocked with normal donkey serum (NDS; 10%, Jackson ImmunoResearch). After the overnight primary antibody incubation at 4°C, the sections were incubated with the secondary antibody (Table 1) for 1 hour at room temperature, followed by a DAPI (1 μg mL^-1^, Sigma-Aldrich) staining for 10 minutes. The sections were mounted with Mowiol (Sigma-Aldrich) and visualized with the Axio Observer 7 microscope (Zeiss).

**Table 1.** The primary and secondary antibodies used for the detection of type I collagen, type II collagen, and SOX9. PBS: phosphate-buffered saline. NGS: normal goat serum. NDS: normal donkey serum.

| **Detection of** | **Primary antibody** | **Secondary antibody** |
|---|---|---|
| Type I collagen | Rabbit anti-collagen I (ab138492; Abcam), 1:400 in PBS with 1% NGS | Goat anti-rabbit Alexa Fluor™ 647 (A21244; Thermo Fisher Scientific), 1:200 in PBS |
| Type II collagen | Mouse anti-collagen II (II-II6B3 supernatant; DSHB), 1:50 in PBS with 1% NGS | Goat anti-mouse Alexa Fluor™ 555 (A21127; Thermo Fisher Scientific), 1:200 in PBS |
| SOX9 | Rabbit anti-SOX9 (PA5-81966; Thermo Fisher Scientific), 1:100 in PBS with 1% NDS | Donkey anti-rabbit Alexa Fluor™<br>647 (711-605-152; Jack-son ImmunoResearch), 1:200 in PBS |
| Ki67 | Rabbit anti-Ki67 (RB-1510-P0; Thermo Fisher Scientific), 1:200 in PBS with 1% NDS | Donkey anti-rabbit Alexa Fluor™<br>647 (711-605-152), 1:200 in PBS |

#### Gene expression analysis

To evaluate the relative gene expression, the tissues were analyzed using quantitative polymerase chain reaction (qPCR). After the culture, the constructs were washed with PBS, snap frozen in liquid nitrogen and stored at −80°C. Frozen samples were crushed using a pestle and treated with RNeasy Lysis Buffer (Qiagen) to lyse the cells. Subsequently, the RNA was isolated with the Qiagen RNeasy kit (Qiagen) following supplier instructions including a 15-minute DNAse (Qiagen) incubation step to remove genomic DNA contamination. After RNA extraction, quantity and purity were assessed with a spectrophotometer (NanoDrop™ One, Isogen Life Science). Next, cDNA was synthesized using Moloney Murine Leukemia Virus Reverse transcriptase (M-MLV RT; 200 U μL^-1^, Invitrogen). Gene expression was evaluated by real-time qPCR (Bio-Rad) using a Fast SYBR green master mix (Applied Biosystems). Here, the qPCR was performed using the primer sequences listed in **Table S1**. The selected primers are coding for proteins related to cell phenotype and matrix deposition. Six reference genes were tested and resulting C_t_ values were normalized for one of the two most stable reference genes, ACTB (β-actin). Relative expression levels were calculated as 2^−ΔCt^ and were compared to those in human neural cells (SH-SY5Y cell line) and human osteoarthritic chondrocytes.

#### Mechanical Testing

The hiCHO-derived organoids and 5w-Dy hiCHO aggregates were evaluated based on their mechanical properties using the MicroTester G2 (CellScale). Unconfined relaxation tests in PBS at room temperature were performed to determine the viscoelastic properties. In these tests, compression until 20% of the diameter of the constructs was applied at 2.5% s^-1^ using a metal plate (2×2 mm) on a cylindrical beam (Ø 0.2032 mm). After compression, the tissue was allowed to relax until equilibrium. Using MATLAB, the stress (*σ)* over time (*t*) was fitted to a standard linear solid (SLS) model (**Equation 1**).^[59,60]^ The contact area between the spherical tissues and metal plate was calculated as described previously.^[25,61]^ With this SLS model, the relaxation time (*τ*) was determined.

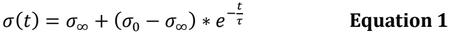

*σ*_∞_= equilibrium stress (Pa)

*σ* _0_= peak stress (Pa)

*t* = time (s)

*τ* = relaxation time (s)

#### Statistical analysis

To evaluate statistical differences between groups, Prism 10.5.0 (GraphPad Software) was used. First, the normality of the data was reviewed using a Shapiro–Wilk test. Data that were not distributed normally or had a small sample size (*N*≤3) were assed using a Mann-Whitney t-test for the comparison of two groups. To compare three or more groups, a Kruskal-Wallis test followed by Dunn’s multiple comparison tests were used. Normally distributed data were assed with a Welch’s t-test or a one-way ANOVA followed by Tukey’s multiple comparison tests. For all statistical tests, P<0.05 was considered significant.

## References

1. Widuchowski W, Widuchowski J, Trzaska T. Articular cartilage defects: Study of 25,124 knee arthroscopies. Knee. 2007;14(3):177–82. doi:10.1016/j.knee.2007.02.001

2. Salzmann GM, Niemeyer P, Hochrein A, Stoddart MJ, Angele P. Articular Cartilage Repair of the Knee in Children and Adolescents. Orthop J Sports Med. 2018;6(3):2325967118760190. doi:10.1177/2325967118760190

3. Merkely G, Ackermann J, Lattermann C. Articular cartilage defects: incidence, diagnosis, and natural history. Oper Tech Sports Med. 2018;26(3):156–61. doi:10.1053/j.otsm.2018.06.008

4. Fox AJS, Bedi A, Rodeo SA. The basic science of articular cartilage: Structure, composition, and function. Sports Health. 2009;1(6):461–8. doi:10.1177/1941738109350438

5. Hunter DJ, Kraus VB. Osteoarthritis — A call to action. Osteoarthritis Cartilage. 2026;34(5):652–3. doi:10.1016/j.joca.2026.02.003

6. Menssen DMA, Vasilopoulou AG, Abinzano F, Korpershoek J V., Kara Özenler A, Papantoniou I, et al. From microtissues to macro solutions –The future of scalable and automated cartilage tissue engineering. J Orthop Translat. 2026;59:101150. doi:10.1016/j.jot.2026.101150

7. Brittberg M, Lindahl A, Nilsson A, Ohlsson C, Isaksson O, Peterson L. Treatment of deep cartilage defects in the knee with autologous chondrocyte transplantation. N Engl J Med. 1994;331(14):889–95. doi:10.1056/NEJM199410063311401

8. Gikas PD, Bayliss L, Bentley G, Briggs TWR. An overview of autologous chondrocyte implantation. J Bone Joint Surg Br. 2009;91(8):997–1006. doi:10.1302/0301-620X.91B8.21824

9. Valisena S, Azogui B, Nizard RS, Tscholl PM, Cavaignac E, Bouché PA, et al. Microfractures, autologous matrix-induced chondrogenesis, osteochondral autograft transplantation and autologous chondrocyte implantation for knee chondral defects: a systematic review and network meta-analysis of randomized controlled trials. EFORT Open Rev. 2024;9(8):785–95. doi:10.1530/EOR-23-0089

10. Hoburg A, Löer I, Körsmeier K, Siebold R, Niemeyer P, Fickert S, et al. Matrix-associated autologous chondrocyte implantation is an effective treatment at midterm follow-up in adolescents and young Adults. Orthop J Sports Med. 2019;7(4):2325967119841077. doi:10.1177/2325967119841077

11. Niemeyer P, Laute V, Zinser W, John T, Becher C, Diehl P, et al. Safety and efficacy of matrix-associated autologous chondrocyte implantation with spheroid technology is independent of spheroid dose after 4 years. Knee Surg Sports Traumatol Arthrosc. 2020;28(4):1130–43. doi:10.1007/s00167-019-05786-8

12. Eschen C, Kaps C, Widuchowski W, Fickert S, Zinser W, Niemeyer P, et al. Clinical outcome is significantly better with spheroid-based autologous chondrocyte implantation manufactured with more stringent cell culture criteria. Osteoarthr Cartil Open. 2020;2(1):100033. doi:10.1016/j.ocarto.2020.100033

13. Textor M, Hoburg A, Lehnigk R, Perka C, Duda GN, Reinke S, et al. Chondrocyte Isolation from Loose Bodies—An Option for Reducing Donor Site Morbidity for Autologous Chondrocyte Implantation. Int J Mol Sci. 2023;24(2):1484. doi:10.3390/ijms24021484

14. Kaps C, Roël G, Eschen C. Method for Cultivation of Cartilage and Spheroids Thereof. Germany; US 2022/0220443 A1, 2022 [cited 2023 Nov 14]. Available from: https://patents.google.com/patent/US20220220443A1/en?oq=US+2022%2f0220443+A1

15. Kisiday JD. Expansion of chondrocytes for cartilage tissue engineering: a review of chondrocyte dedifferentiation and redifferentiation as a function of growth in expansion culture. Regen Med Front. 2020;2(1):e200002. doi:10.20900/RMF20200002

16. Su W, Nie Y, Zheng S, Yao Y. Recent Research on Chondrocyte Dedifferentiation and Insights for Regenerative Medicine. Biotechnol Bioeng. 2025;122(4):749–60. doi:10.1002/bit.28915

17. McCarthy HS, Tins B, Gallacher PD, Jermin P, Richardson JB, Kuiper JH, et al. Histological and Radiological Assessment of Endogenously Generated Repair Tissue In Vivo Following a Chondral Harvest. Cartilage. 2023;14(1):48–58. doi:10.1177/19476035221149523

18. Mazzini G, Houtman E, Hoolwerff M van, Janssen M, Sayedipour S, Hajmousa G, et al. Dynamic suspension culture enhances scalable maturation of hiPSC-derived cartilage organoids for regenerative medicine. bioRxiv. 2026. doi:10.64898/2026.06.15.732328

19. Abe K, Yamashita A, Morioka M, Horike N, Takei Y, Koyamatsu S, et al. Engraftment of allogeneic iPS cell-derived cartilage organoid in a primate model of articular cartilage defect. Nat Commun. 2023;14(1):804. doi:10.1038/s41467-023-36408-0

20. Yamashita A, Morioka M, Yahara Y, Okada M, Kobayashi T, Kuriyama S, et al. Generation of scaffoldless hyaline cartilaginous tissue from human iPSCs. Stem Cell Reports. 2015;4(3):404–18. doi:10.1016/j.stemcr.2015.01.016

21. Fujisawa Y, Takao T, Yamada D, Takarada T. Development of cartilage tissue using a stirred bioreactor and human iPSC-derived limb bud mesenchymal cells. Biochem Biophys Res Commun. 2023;687:149146. doi:10.1016/j.bbrc.2023.149146

22. Rodríguez Ruiz A, Dicks A, Tuerlings M, Schepers K, van Pel M, Nelissen RGHH, et al. Cartilage from human-induced pluripotent stem cells: comparison with neo-cartilage from chondrocytes and bone marrow mesenchymal stromal cells. Cell Tissue Res. 2021;386(2):309–20. doi:10.1007/s00441-021-03498-5

23. Crispim JF, Ito K. De novo neo-hyaline-cartilage from bovine organoids in viscoelastic hydrogels. Acta Biomater. 2021;128:236–49. doi:10.1016/j.actbio.2021.04.008

24. Kleuskens MWA, Crispim JF, van Donkelaar CC, Janssen RPA, Ito K. Evaluating initial integration of cell-based chondrogenic constructs in human osteochondral explants. Tissue Eng Part C Methods. 2022;28(1):34–44. doi:10.1089/ten.tec.2021.0196

25. Menssen DMA, Feenstra JCA, Janssen RPA, Abinzano F, Ito K. Cartilage organoids from articular chondroprogenitor cells and their potential to produce neo-hyaline cartilage. Cartilage. 2025;0(0):19476035241313179. doi:10.1177/19476035241313179

26. Hajmousa G, de Almeida RC, Bloks N, Ruiz AR, Bouma M, Slieker R, et al. The role of DNA methylation in chondrogenesis of human iPSCs as a stable marker of cartilage quality. Clin Epigenetics. 2024;16(1):141. doi:10.1186/s13148-024-01759-y

27. Kamaraj A, Kyriacou H, Seah KTM, Khan WS. Use of human induced pluripotent stem cells for cartilage regeneration in vitro and within chondral defect models of knee joint cartilage in vivo: a Preferred Reporting Items for Systematic Reviews and Meta-Analyses systematic literature review. Cytotherapy. 2021;23(8):647–61. doi:10.1016/j.jcyt.2021.03.008

28. Yoshihara M, Hayashizaki Y, Murakawa Y. Genomic instability of iPSCs: challenges towards their clinical applications. Stem Cell Rev Rep. 2017;13(1):7–16. doi:10.1007/s12015-016-9680-6

29. Mulders R, Nickel-Maunu M, van Hoolwerff M, Mazzini G, Klomp L, Meijer H, et al. Nuclear Factor I genes drive chondrogenic cell-fate commitment. bioRxiv. 2026;719911. doi:10.64898/2026.04.21.719911

30. Pfister P, Lhospice E, García-García A, Paillaud R, Jung S, Schaller R, et al. Start, Stop, Rewind, Repeat—Cyclic Exposure of Adipose Stromal Cells-derived Cartilage Organoids to Chondrogenic and Proliferative Cues to Achieve Scaled-up and Customizable Bone Formation by Endochondral Ossification. Adv Healthc Mater. 2026. doi:10.1002/adhm.202504880

31. Büttner P, Faber J, Teßmar J, Stahlhut P, Budday S, Blunk T. Tissue maturation and development of mechanical properties in hyaluronic acid bioink-based cartilaginous constructs. Front Bioeng Biotechnol. 2026;14:1749259. doi:10.3389/fbioe.2026.1749259

32. Wong M, Ponticiello M, Kovanen V, Jurvelin JS. Volumetric changes of articular cartilage during stress relaxation in unconfined compression. J Biomech. 2000;33(9):1049–54. doi:10.1016/S0021-9290(00)00084-1

33. Lau YT, Li HC, Ho WYV, Chan BP. Modular assembly of human auricular chondrocyte-derived microtissues for ear cartilage tissue engineering. Acta Biomater. 2026. doi:10.1016/j.actbio.2026.07.019

34. Nakamura A, Zujur D, Nonaka T, Yoshizato H, Kashimoto S, Ikeya M, et al. Modulation of iPSC-derived NCMSC cell state by TD-198946 enhances scaffold-free cartilage biofabrication. Biofabrication. 2026. doi:10.1088/1758-5090/ae44ed

35. Parreno J, Raju S, Wu P han, Kandel RA. MRTF-A signaling regulates the acquisition of the contractile phenotype in dedifferentiated chondrocytes. Matrix Biology. 2017;62:3–14. doi:10.1016/j.matbio.2016.10.004

36. Yasui N, Nimni ME. Cartilage Collagens. In: Collagen, Volume I: Biochemistry. 1st Edition. Boca Raton; 1988. p. 225–41. doi:10.1201/9781351070799

37. Li H, Haudenschild DR, Posey KL, Hecht JT, Di Cesare PE, Yik JHN. Comparative analysis with collagen type II distinguishes cartilage oligomeric matrix protein as a primary TGFβ-responsive gene. Osteoarthritis Cartilage. 2011;19(10):1246–53. doi:10.1016/j.joca.2011.07.011

38. Shepherd C, Reese AE, Reynard LN, Loughlin J. Expression analysis of the osteoarthritis genetic susceptibility mapping to the matrix Gla protein gene MGP. Arthritis Res Ther. 2019;21(1):149. doi:10.1186/s13075-019-1934-7

39. Wu CL, Dicks A, Steward N, Tang R, Katz DB, Choi YR, et al. Single cell transcriptomic analysis of human pluripotent stem cell chondrogenesis. Nat Commun. 2021;12(1):362. doi:10.1038/s41467-020-20598-y

40. Dicks A, Wu CL, Steward N, Adkar SS, Gersbach CA, Guilak F. Prospective isolation of chondroprogenitors from human iPSCs based on cell surface markers identified using a CRISPR-Cas9-generated reporter. Stem Cell Res Ther. 2020;11(1):66. doi:10.1186/s13287-020-01597-8

41. Adkar SS, Wu CL, Willard VP, Dicks A, Ettyreddy A, Steward N, et al. Step-wise chondrogenesis of human induced pluripotent stem cells and purification via a reporter allele generated by CRISPR-Cas9 genome editing. Stem Cells. 2019;37(1):65–76. doi:10.1002/stem.2931

42. Niemeyer P, Laute V, John T, Becher C, Diehl P, Kolombe T, et al. The Effect of Cell Dose on the Early Magnetic Resonance Morphological Outcomes of Autologous Cell Implantation for Articular Cartilage Defects in the Knee. American Journal of Sports Medicine. 2016;44(8):2005–14. doi:10.1177/0363546516646092

43. Burdis R, Chariyev-Prinz F, Browe DC, Freeman FE, Nulty J, McDonnell EE, et al. Spatial patterning of phenotypically distinct microtissues to engineer osteochondral grafts for biological joint resurfacing. Biomaterials. 2022;289:121750. doi:10.1016/j.biomaterials.2022.121750

44. de Vries S, Doeselaar M van, Meij B, Tryfonidou M, Ito K. Notochordal Cell Matrix As a Therapeutic Agent for Intervertebral Disc Regeneration. Tissue Eng Part A. 2019;25(11–12):830–41. doi:10.1089/ten.tea.2018.0026

45. de Vries SAH, Potier E, van Doeselaar M, Meij BP, Tryfonidou MA, Ito K. Conditioned Medium Derived from Notochordal Cell-Rich Nucleus Pulposus Tissue Stimulates Matrix Production by Canine Nucleus Pulposus Cells and Bone Marrow-Derived Stromal Cells. Tissue Eng Part A. 2015;21(5–6):1077–84. doi:10.1089/ten.tea.2014.0309

46. Bach F, Libregts S, Creemers L, Meij B, Ito K, Wauben M, et al. Notochordal-cell derived extracellular vesicles exert regenerative effects on canine and human nucleus pulposus cells. Oncotarget. 2017;8(51):88845–56. doi:10.18632/oncotarget.21483

47. Müller S, Acevedo L, Wang X, Karim MZ, Matta A, Mehrkens A, et al. Notochordal cell conditioned medium (NCCM) regenerates end-stage human osteoarthritic articular chondrocytes and promotes a healthy phenotype. Arthritis Res Ther. 2016;18(1):125. doi:10.1186/s13075-016-1026-x

48. Ansari M, Darvishi A, Sabzevari A. A review of advanced hydrogels for cartilage tissue engineering. Front Bioeng Biotechnol. 2024;12:1340893. doi:10.3389/fbioe.2024.1340893

49. van de Looij SM, Vasilopoulou AG, Spauwen L, van den Dikkenberg A, Korpershoek J V., de Ruijter M, et al. Diels-alder click chemistry as a dynamic-covalent crosslinking method in spheroid-encapsulating hydrogels for cartilage engineering. Adv Healthc Mater. 2026;e05013. doi:10.1002/adhm.202505013

50. Dufour A, Gallostra XB, O’Keeffe C, Eichholz K, Von Euw S, Garcia O, et al. Integrating melt electrowriting and inkjet bioprinting for engineering structurally organized articular cartilage. Biomaterials. 2022;283:121405. doi:10.1016/j.biomaterials.2022.121405

51. Spauwen L, Pueyo Moliner A, van Veenendaal P, Custers R, Malda J, de Ruijter M. Engineering anisotropic mechanical properties in large-scale fabricated cartilage constructs using microfiber reinforcement. Adv Healthc Mater. 2025;14(19):2501014. doi:10.1002/adhm.202501014

52. Peters JR, Hoogenboom M, Abinzano F, Callens SJP, Foolen J, Ito K. Tissue growth as a mechanism for collagen fiber alignment in articular cartilage. Sci Rep. 2024;14(1):31121. doi:10.1038/s41598-024-82441-4

53. Licata JP, Schwab KH, Har-el Y el, Gerstenhaber JA, Lelkes PI. Bioreactor technologies for enhanced organoid culture. Int J Mol Sci. 2023;24(14):11427. doi:10.3390/ijms241411427

54. Das R, Roosloot R, van Pel M, Schepers K, Driessen M, Fibbe WE, et al. Preparing for cell culture scale-out: establishing parity of bioreactor- and flask-expanded mesenchymal stromal cell cultures. J Transl Med. 2019;17(1):241. doi:10.1186/s12967-019-1989-x

55. Dambrot C, van de Pas S, van Zijl L, Brändl B, Wang JW, Schalij MJ, et al. Polycistronic lentivirus induced pluripotent stem cells from skin biopsies after long term storage, blood outgrowth endothelial cells and cells from milk teeth. Differentiation. 2013;85(3):101–9. doi:10.1016/j.diff.2013.01.001

56. Bomer N, den Hollander W, Suchiman H, Houtman E, Slieker RC, Heijmans BT, et al. Neo-cartilage engineered from primary chondrocytes is epigenetically similar to autologous cartilage, in contrast to using mesenchymal stem cells. Osteoarthr Cartil. 2016;24(8):1423–30. doi:10.1016/j.joca.2016.03.009

57. de Vries SAH, van Doeselaar M, Kaper HJ, Sharma PK, Ito K. Notochordal cell matrix as a bioactive lubricant for the osteoarthritic joint. Sci Rep. 2018;8(1):8875. doi:10.1038/s41598-018-27130-9

58. Farndale RW, Sayers CA, Barrett AJ. A Direct Spectrophotometric Microassay for Sulfated Glycosaminoglycans in Cartilage Cultures. Connect Tissue Res. 1982;9(4):247–8. doi:10.3109/03008208209160269

59. Richard F, Villars M, Thibaud S. Viscoelastic modeling and quantitative experimental characterization of normal and osteoarthritic human articular cartilage using indentation. J Mech Behav Biomed Mater. 2013;24:41–52. doi:10.1016/j.jmbbm.2013.04.012

60. Lin CY. Rethinking and researching the physical meaning of the standard linear solid model in viscoelasticity. Mechanics of Advanced Materials and Structures. 2024;31(11):2370–85. doi:10.1080/15376494.2022.2156638

61. Kim K, Cheng J, Liu Q, Wu XY, Sun Y. Investigation of mechanical properties of soft hydrogel microcapsules in relation to protein delivery using a MEMS force sensor. J Biomed Mater Res A. 2010;92A(1):103–13. doi:10.1002/jbm.a.32338

