## Supplementary material for "Extracellular matrix-enriched human induced chondrocyte organoids as living building blocks for scalable cartilage tissue engineering": Table S1

### **Table S1. Primer sequences of the analyzed genes.**

| **Gene** | **Full gene name** | **Role** | **5′–3′** |
| --- | --- | --- | --- |
| *MGP* | Matrix Gla Protein | Hypertrophy | F - CCGCCTTAGCGGTAGTAACTT |
|  |  |  | R - GAGCGTTCTCGGATCCTCTC |
| *COMP* | Cartilage Oligomeric Matrix Protein | ECM organization | F - GGGTTACACTGCCTTCAATGG |
|  |  |  | R - AGCTGGAGCTGTCCTGGTAG |
| *CHAD* | Chondroadherin | Adhesion | F - GGCCGCCATTAAACAGGTTC |
|  |  |  | R - AGGCTGTGGGGAGAAGGTAT |
| *SOX9* | SRY-Box Transcription Factor 9 | Chondrogenesis | F -GGCAAGCTCTGGAGACTTCTG |
|  |  |  | R -CCCGTTCTTCACCGACTTCC |
| *ACAN* | Aggrecan | Cartilage matrix proteoglycan | F -TGCTTTGTAGACAGACTTGAGG |
|  |  |  | R -CCAGCGTAGCATTGTGAGATT |
| *COL1A1* | Collagen Type I Alpha 1 Chain | Fibrocartilage | F -AATCACCTGCGTACAGAACGG |
|  |  |  | R -TCGTCACAGATCACGTCATCG |
| *COL2A1* | Collagen Type II Alpha 1 Chain | Cartilage matrix protein | F -GGGAGAGCCTGGAGATGAC |
|  |  |  | R -GACCGACGATGCCTCTCTG |
| *COL10A1* | Collagen Type X Alpha 1 Chain | Hypertrophy | F -GTGGACCAGGAGTACCTTGC |
|  |  |  | R -CATAAAAGGCCCACTACCCA |
| *KIF1A* | Kinesin Family Member 1A | Neurogenic | F -CGGGAAATGAGCCGTGACT |
|  |  |  | R -CGACGCGTAGTTGATGTCCT |
| *ACTB* | Actin Beta | Reference gene | F - ACAGAGCCTCGCCTTTGCC |
|  |  |  | R - GAGGATGCCTCTCTTGCTCTG |
