## Supplementary material for "Extracellular matrix-enriched human induced chondrocyte organoids as living building blocks for scalable cartilage tissue engineering": Figure S1

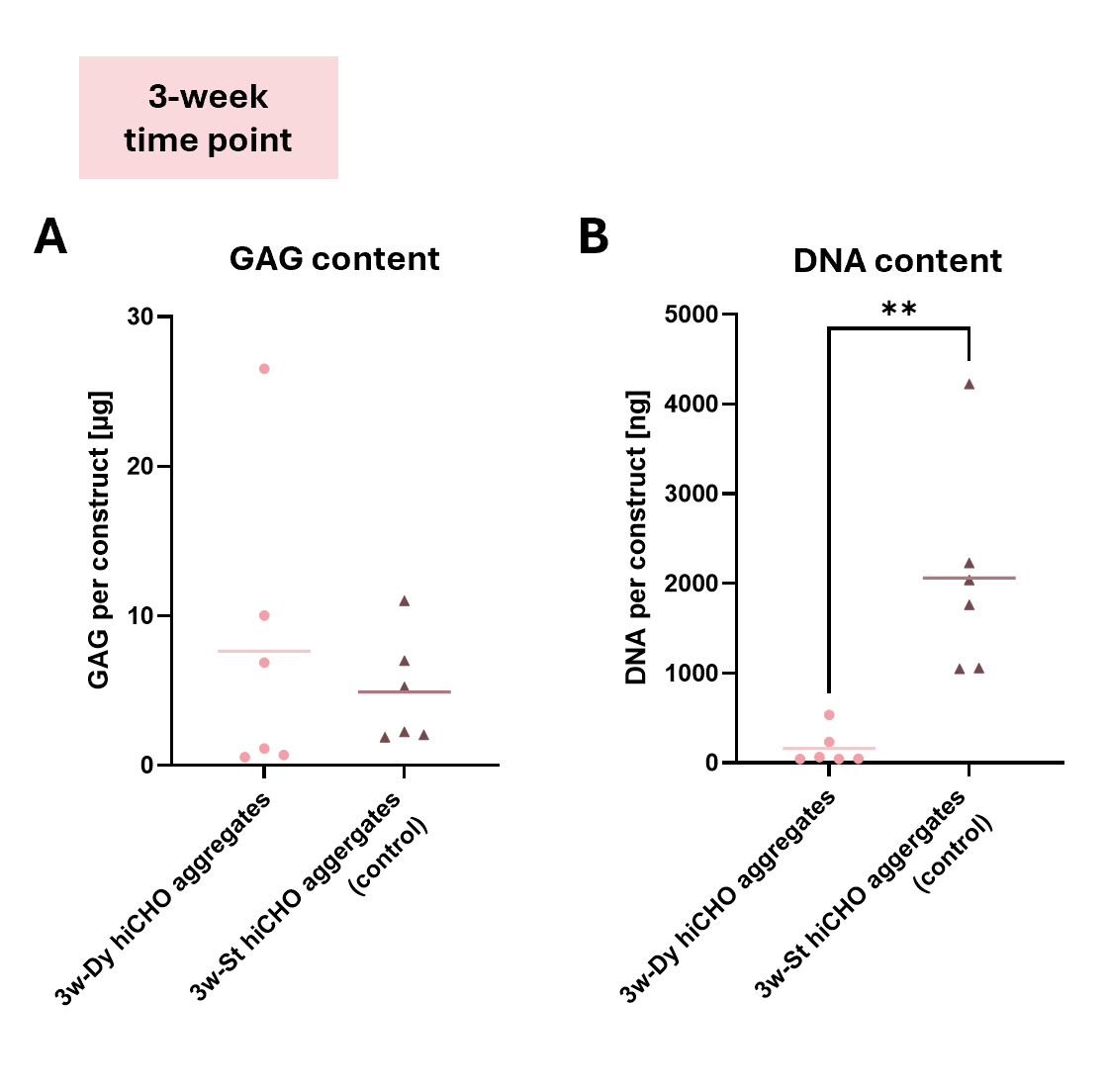

Figure S1. Quantification of sulfated GAGs per 3w-Dy hiCHO aggregate and 3w-St hiCHO aggregate (A) and of DNA per 3w-Dy hiCHO aggregate and 3w-St hiCHO aggregate (B). The results of the Mann-Whitney t-tests are depicted in the graphs, where ** indicates P<0.01.
